# Identification of a subpopulation of highly adherent endothelial cells for seeding synthetic vascular grafts

**DOI:** 10.1101/2023.08.25.554908

**Authors:** Jayne T. Wolfe, Vaya Chen, Yiliang Chen, Brandon J. Tefft

## Abstract

**Objective:** There is an unmet clinical need for a bypass graft that can be used as an alternative to an autologous vessel graft for the treatment of severe coronary artery disease. Small-diameter (<6mm) synthetic vascular grafts are not suitable because of unacceptable patency rates. This mainly occurs without an endothelial cell (EC) monolayer to prevent platelet activation, thrombosis, and intimal hyperplasia. While numerous studies have explored methods to improve EC adhesion to biomaterials, there are still no reliable methods to endothelialize small-diameter grafts, as most seeded ECs are lost due to exposure to fluid shear stress (SS) after implantation. The goal of this work is to determine if EC loss is a random process or if it is possible to predict which cells are more likely to remain adherent.

**Approach and Results:** In initial studies, we sorted ECs using fluid SS and identified a subpopulation of ECs that are more likely to resist detachment. We use RNA-sequencing (RNA-seq) to examine gene expression of adherent ECs compared to the whole population to identify targets for improving adhesion. Fibronectin leucine rich transmembrane protein 2 (*FLRT2*), which encodes protein FLRT2, emerged as a candidate due to its downregulation in the adherent ECs and known role in cell adhesion. Using fluorescence activated cell sorting (FACS), we sorted ECs based on FLRT2 expression levels and demonstrated that ECs expressing low levels of FLRT2 exhibit greater retention under fluid SS in vitro.

**Conclusion:** For the first time, we show EC detachment is not an entirely random process and we predicted which ECs were more likely to remain adherent on a vascular graft upon exposure to fluid SS. This provides validation for the concept that we can seed a small-diameter vascular graft only with highly adherent ECs to maintain a stable endothelium and improve graft patency rates.

**Non-standard Abbreviations and Acronyms:** endothelial cell (EC), shear stress (SS), fibronectin leucine rich transmembrane protein 2 (FLRT2), tissue engineered vascular graft (TEVG), fluorescence activated cell sorting (FACS)

## Introduction

Patients with severe coronary artery disease often require surgical intervention with a bypass graft to restore blood flow. Autologous vessels remain the only effective graft option, yet more than 30% of patients with cardiovascular disease do not have suitable vasculature that can be used for autologous bypass grafting.^1^ Early reports using synthetic vascular grafts described 64% of 4mm diameter expanded polytetrafluorethylene (ePTFE) grafts were patent in patients 1 year after surgery and only 14% remained patent after 3 years.^2^ Therefore, there is an urgent unmet clinical need to improve the patency rates of small-diameter synthetic vascular grafts (<6mm) to overcome the limitations of autologous vessel grafts for applications including coronary artery bypass graft (CABG) procedures.

All blood-contacting devices that lack a stable endothelial cell (EC) monolayer are vulnerable to thrombosis after implantation. In the native blood vessel, ECs form a monolayer on the vessel wall and modulate the blood-tissue interface to inhibit platelet aggregation.^3^ All implanted biomaterials exhibit some degree of thrombogenicity because of nonspecific adsorption of serum proteins to the biomaterial surface, which then bind to circulating platelets. This leads to platelet activation, aggregation, and subsequent thrombosis, resulting in loss of patency.^4^ Neointimal hyperplasia can also lead to loss of graft patency as smooth muscle cells proliferate and migrate into the sub-endothelial space, resulting in thickening of the vessel wall.^5^ ECs regulate neointimal growth by inhibiting excessive smooth muscle cell proliferation and migration, especially at the anastomoses.^6^

Tissue engineered vascular grafts (TEVGs) containing a living endothelium have been proposed to achieve acceptable long-term patency.^7-9^ Despite numerous attempts to endothelialize synthetic vascular grafts, most seeded ECs are lost from the biomaterial surface due to exposure to fluid shear stress (SS) from circulating blood.^10^ The mechanism driving some ECs to detach while others remain adherent is still unknown. It may be driven by lack of mature focal adhesions.^11^ The goal of this work is to investigate the molecular mechanisms that permit some ECs to resist detachment upon exposure to fluid SS and to use this knowledge to enrich for the ECs that are most likely to resist detachment. Ultimately, this approach can be applied to fabricate vascular grafts containing a stable EC monolayer.

In this study, we identify a subpopulation of ECs that are more likely to resist detachment upon exposure to fluid SS *in vitro*. While the response of ECs to SS has been thoroughly examined in the literature,^12^ we use SS as a means to sort an EC population into subpopulations capable of resisting detachment for increasing durations of time. We hypothesize that differential expression of genes involved in cell adhesion pathways allows the ECs in the adherent subpopulations to remain adherent. We use bulk RNA-sequencing (RNA-seq) to compare the transcriptome of adherent subpopulations of ECs compared to the whole population of ECs. We show that fibronectin leucine-rich transmembrane protein 2 (FLRT2) can be used as a marker to select for the highly adherent ECs using fluorescence activated cell sorting (FACS). The sorted ECs expressing low levels of FLRT2 demonstrated enhanced retention on both tissue culture plates and vascular grafts when exposed to fluid SS.

## Methods

### Cell Culture

For the initial investigations of EC retention, porcine blood outgrowth endothelial cells (BOECs) were used. Our lab has an established porcine model for conducting vascular graft studies *in vivo* and autologous BOECs can be generated for the animals using peripheral blood. BOECs were generated from peripheral porcine blood as previously described.^13^ BOECs were maintained in endothelial growth medium-2 (EGM-2) (#CC-3162, Lonza, Portsmouth, NH) supplemented with 10% fetal bovine serum (#F08BB22A1, Atlas Biologicals, Fort Collins, CO) and 1× antibiotic-antimycotic (Fisher Scientific, Hampton, MA). For investigations of EC transcriptome, primary human umbilical vein endothelial cells (HUVECs) were used (#PCS-100-013, ATCC, Manassas, VA). This cell type provides a model for studying human gene expression, which is more relevant for future clinical applications. HUVECs are a widely used cell type for studying EC adhesion and the response of ECs to SS.^14^ These cells have also been used in RNA-sequencing studies and would allow us to compare our data to other available RNA-seq datasets.^15^ For all other experiments, pooled primary HUVECs (#PCS-100-014, ATCC, Manassas, VA) were used to avoid subject-specific variability. HUVECs were maintained in vascular cell basal medium (#PCS-100-030, ATCC, Manassas, VA) supplemented with the endothelial cell growth kit bovine brain extract (BBE) (#PCS-100-040, ATCC, Manassas, VA), 10% fetal bovine serum, and 1× antibiotic-antimycotic (ThermoFisher Scientific, Waltham, MA). Passage 3-5 cells were used for experiments.

### Retention Experiments Using Shear Stress and Trypsin Detachment

For SS isolation experiments, BOECs were cultured overnight on fibronectin-coated (0.2 mg/mL) 100 mm dishes in the flow path area using a rectangular gasket (#31-011, Glycotech, Gaithersburg, MD). A rectangular parallel plate flow chamber (#31-010, GlycoTech, Gaithersburg, MD) was placed on top of the cultured cells using vacuum pressure to form a seal. A multi-channel syringe pump (NE-1600, New Era Pump Systems, Farmingdale, NY) was used to infuse up to six parallel plate flow chambers at once. A flow rate of 3.18 mL/min was used to achieve a fluid SS of 15 dyn/cm^2^ at the cell surface based on the gasket and dimensions and fluid mechanics theory describing flow between parallel plates. BOECs were exposed to fluid SS for 30 min. Non-adherent ECs that detached were discarded. Adherent BOECs in the flow path area were collected using 3 min incubation with 0.25% trypsin/EDTA (#25200056, Fisher Scientific, Hampton, MA) and centrifuged at 300×g for 5 min to obtain a cell pellet. Adherent BOECs were re-plated and allowed to adhere for 3 hrs. Control samples were plated using the whole population ECs and allowed to adhere for 3 hrs. Cell nuclei were stained with Hoechst solution (#33342, ThermoFisher Scientific, Waltham, MA) and fluorescence microscopy was used to image the cells (NS-100, Nikon, Melville, NY). Each plate was exposed to 15 dyn/cm^2^ fluid SS for 30 min again as described above. Images obtained after exposure to fluid SS were compared to baseline images to calculate cell retention percentage. A total of n=5-11 plates per group were used.

For trypsin isolation experiments, a T-75 flask of BOECs was incubated with 0.25% trypsin/EDTA solution (#25200056, Fisher Scientific, Hampton, MA) for 3 min until approximately 80% of BOECs were detached. These BOECs were collected and designated as non-adherent. The remaining adherent BOECs were incubated an additional 1 min with trypsin until all cells detached from the flask. These BOECs were collected and designated as adherent. For the whole population, another T-75 flask was incubated with trypsin for 4 min to obtain the whole population with both the adherent and non-adherent subpopulations. All populations were centrifuged at 300×g for 5 min to obtain a cell pellet and re-plated on fibronectin-coated tissue culture plates. The cell populations were allowed to adhere overnight and then subjected to 30 min of 15 dyn/cm^2^ fluid SS using a parallel plate flow chamber to measure cell retention as described above. A total of n=6 plates per group were used.

### Shear Stress Experiments for RNA-sequencing

Primary HUVECs were cultured overnight on fibronectin-coated 100 mm dishes in the flow path area using a rectangular gasket and then subjected to 15 dyn/cm^2^ fluid SS using a rectangular parallel pate flow chamber as described above. HUVECs were exposed to SS for 0, 5, 10, or 30 min to generate the whole population, 5 min adherent, 10 min adherent, and 30 min adherent subpopulations, respectively. Plates from each time point were pooled together to generate sufficient RNA for analysis. A total of n=3 replicates from independent experiments were used for each time point.

### RNA Extraction

Immediately after exposure to fluid SS, retained HUVECs were washed once with phosphate buffered saline (PBS) (non-retained HUVECs were discarded). Cells were incubated with 0.25% trypsin/EDTA (#25200056, Fisher Scientific, Hampton, MA) for 3 min and centrifuged at 300×g for 5 min to collect in a cell pellet. RNA was extracted and purified according to the RNeasy Plus Mini Kit protocol (#74134, Qiagen, Germantown, MD) with QIA shredder homogenization columns (#79654, Qiagen, Germantown, MD) and gDNA elimination columns (#74134, Qiagen, Germantown, MD n). RNA concentration and quality were assessed using a NanoDrop (#A38189, ThermoFisher Scientific, Waltham, MA). Samples were stored in RNase free H_2_O (#74134, Qiagen, Germantown, MD) at -80°C until sent for sequencing.

### RNA-sequencing

RNA-sequencing was performed by ArrayStar (Rockville, MD). Total RNA was enriched by oligo (dT) magnetic beads to remove rRNA. RNA-seq library preparation was performed using KAPA Stranded RNA-Seq Library Prep Kit (Illumina, San Diego, CA). Completed libraries were qualified (Aligent 2100 Bioanalyzer) and quantified (qPCR for absolute quantification). Libraries were sequenced on the NovaSeq 6000 (Illumina, San Diego, CA). Sequence quality was verified using FastQC software.^16^ Trimmed reads (trimmed 5’, 3-adaptor bases with cutadapt^17^) were aligned to the human reference genome with Hisat2 software.^18^ Transcript abundances were estimated using StringTie.^19^ Fragments per kilobase of transcript per million mapped reads (FPKM) values^20^ for expressed genes and transcripts were filtered with Ballgown R package.^20-22^

### Immunofluorescence

Whole population (0 min fluid SS) and adherent subpopulation (30 min fluid SS) plates of HUVECs were exposed to 0 and 30 min of 15 dyn/cm^2^ fluid SS, respectively, using a circular parallel plate flow chamber as described above. Adherent ECs were fixed with 4% paraformaldehyde and permeabilized using 0.2% Triton-X in PBS. Cells were blocked in goat serum for 1 hr (#31872, Fisher Scientific, Hampton, MA). Cells were incubated with vinculin primary antibody (#V9264, MilliPore Sigma, Burlington, MA) at 10 µg/mL in PBS for 1 hr at 37°C. The primary antibody was removed, and cells were washed 3× with PBS. Goat anti-mouse Alexa Fluor 488 (#A-11001, ThermoFisher Scientific, Waltham, MA) secondary antibody and DAPI (#62248, ThermoFisher Scientific, Waltham, MA) were added at 1:750 in PBS. Cells were incubated for 1 hr at 37°C. Cells were washed with PBS 3× then covered in PBS for imaging with a fluorescence microscope (NS-100, Nikon, Melville, NY).

### Focal Adhesion Quantification

The total number and area of vinculin-containing focal adhesions (FAs) were quantified from the images with vinculin staining in ImageJ Fiji^23^ according to methods previously described by Horzum et al.^24^ First, “subtract background” was applied with the sliding paraboloid option with the radius set to 50 pixels. The local contrast was enhanced using the CLAH plugin. The Brightness & Contrast was adjusted automatically. The mathematical EXP command was used to minimize the background. The Mexican Hat filter plugin was used with radius set to 8. The Threshold command was used to convert the grayscale image to binary using the automatic setting. The number of FAs and FA area were measured using the binary image and Analyze Particles command. The Analyze Particles was set to circularity=0.40-0.99 and show masks. A total of 3-5 images from each of the n=3 plates per group were used for analysis.

### Quantitative RT-qPCR

Extracted total RNA was reverse transcribed into cDNA using the QuantiTect Reverse Transcription Kit protocol (#205311, Qiagen, Germantown, MD). Pre-designed primers were purchased for FLRT2 (#Hs00544171_s1, ThermoFisher Scientific, Waltham, MA) and housekeeping gene for beta tubulin (TUBB) (#Hs00742828_s1, ThermoFisher Scientific, Waltham, MA). TaqMan master mix (4444556, ThermoFisher Scientific, Waltham, MA) and 384-well plates (#<u>A36931</u>, ThermoFisher Scientific, Waltham, MA) were used with the RT-qPCR machine (QuantStudio 6 Pro Applied Biosystems, ThermoFisher Scientific, Waltham, MA). The expression of *FLRT2* was determined relative to *TUBB* using the delta-delta CT method. A total of n=6 replicates per sample were used.

### Flow Cytometry

Pooled primary HUVECs were collected using incubation with Accutase cell detachment solution (#SCR005, MilliporeSigma, Burlington MA) for 10 min at 37°C. Cells were centrifuged at 300×g for 5 min and washed with 10 mL cell staining buffer (#420201, Biolegend, San Diego, CA). Human FLRT2 antibody (#AF2877, Invitrogen R&D Systems, Minneapolis, MN) was added at 20 μg/mL in 100 μl stain buffer per 1x10^6^ cells and incubated for 20 min at room temperature protected from light. For the IgG control, goat IgG (#AB-108-C, Invitrogen R&D Systems, Minneapolis, MN) was added at 20 μg/mL in 100 μl stain buffer per 1×10^6^ cells and incubated for 20 min at room temperature protected from light. Cells were washed with 4mL cell staining buffer and pelleted with centrifugation at 300×g for 5 min. Donkey anti-goat Alexa Fluor 488 (#A11055, ThermoFisher Scientific, Waltham, MA) was added at 2 μg/mL in 100 μl stain buffer per 1×10^6^ cells and incubated for 20 min at room temperature protected from light. Cells were washed with 4mL cell staining buffer and pelleted with centrifugation at 300×g for 5 min. Cells were resuspended in 1mL cell staining buffer (sterile filtered with 0.2μm filter) and analyzed using a flow cytometer (LSRII, Becton Dickinson, Vernon Hills, IL). Mean fluorescence intensity of Alexa Fluor 488 was calculated using FlowJo software (Becton Dickinson, Vernon Hills, IL). A total of n=5 independent experiments were performed.

### Fluorescence Activated Cell Sorting (FACS)

Pooled primary HUVECs were prepared for FACS using the flow cytometry protocol described above. FACS was performed using a cell sorter (Aria, Becton Dickinson, Vernon Hills, IL). Gates were applied to separate cells into low, intermediate, and high staining groups. The unstained control sample was used to set the gate for the unstained cells in the “low” expression group. The positively stained cells were divided approximately in half to set the gates to obtain the “intermediate” and “high” expression groups. The same gates were applied for all sorting experiments. Cells were collected in the HUVEC media described above supplemented with 10 μg/mL gentamicin (#15710064, ThermoFisher Scientific, Waltham, MA).

### Sorted Endothelial Cell Retention Experiments on Tissue Culture Plates

Pooled primary HUVECs were sorted using FACS into low, intermediate, and high FLRT2 expression groups. Sorted HUVECs were seeded onto fibronectin-coated 35 mm dishes in the circular chamber flow path area and cultured overnight. Cell retention after exposure to 30 min of 15 dyn/cm^2^ fluid SS with a circular parallel plate flow chamber was quantified with nuclear imaging as described above. A total of n=5-6 plates per group were tested.

### Sorted Endothelial Cell Retention Experiments on Vascular Grafts

Polyester urethane (DEGRAPOL® DP15, Ab Medica S.p.A.) 4 mm inner-diameter vascular grafts were fabricated by electrospinning (SprayBase, Avectas). A 20% wt/v (DP15/chloroform) solution was electrospun using 0.2barr pressure, ∼11 kV applied voltage, 15 cm emitter to mandrel distance, 4 mm diameter mandrel, and 450 rpm mandrel rotation speed. Samples with 4 mm inner diameter, 0.8 mm wall thickness, and 3-4 cm length were fabricated for retention studies. Samples were sterilized using ethylene oxide. Pooled primary HUVECs were sorted using FACS into low and FLRT2 expression groups. The same gate for the low expression group was chosen based off the unstained control sample. The high gate included the positively stained cells previously defined in the both the intermediate and high groups for the retention studies on the tissue culture plates. Sorted HUVECs were seeded onto fibronectin-coated vascular graft samples at a density of 200,000 cells/cm^2^ and cultured overnight. Grafts were mounted in a custom bioreactor chamber (Engineering Core, Medical College of Wisconsin) and exposed to a mean fluid SS of 10 dyn/cm^2^ via warmed PBS perfused by a pulsatile blood pump (#55-1838, Harvard Apparatus, Holliston, MA). The fluid SS value of 10 dyn/cm^2^ was chosen based on the average SS in the coronary artery, which is reported to be 7 dyn/cm^2^. Initially we used 15 dyn/cm^2^ fluid SS to identify the most adherent ECs in the population and then switched to 10 dyn/cm^2^ fluid SS to subject the sorted ECs to a more physiologic fluid SS level. The pump was set to 10 mL stroke volume and 42 strokes/min to achieve a mean flow rate of 420 mL/min. Following exposure to 30 min of fluid SS, grafts were fixed with 4% paraformaldehyde and washed 3× with PBS. Nuclei were stained with DAPI (#R37606, ThermoFisher Scientific, Waltham, MA) and grafts were sectioned open lengthwise. Grafts were mounted on a slide with coverglass and imaged *en face* using a fluorescence microscope as described above. To calculate the cell retention, a seeding control was used for each sample since the same graft could not be imaged before and after exposure to fluid SS. The same cell population was seeded onto the seeding control at the time of seeding the experimental grafts for each experiment. The cell number after exposure to fluid SS was compared to the cell number from the seeding control not exposed to fluid SS and cell retention percentage was calculated. A total of n=5 grafts per group were tested.

### Statistical Analysis

A t-test was used to compare the means of two groups and a one-way analysis of variance (ANOVA) with Tukey post-hoc analysis was used to compare the means of three or more groups. Statistical significance was defined as p<0.05. All results are reported as the mean ± standard error of the mean.

## Results

### Endothelial cell detachment is not an entirely random process

Flow chamber experiments identified a subpopulation of adherent ECs that resists detachment when exposed to fluid SS and provided evidence that EC detachment is not an entirely random process (Figure 1). We used fluid SS to obtain a population of ECs that resist detachment. ECs were subjected to 15 dyn/cm^2^ of fluid SS for 30 min. This SS value was chosen based on the upper limit of the average wall SS values measured or predicted in the human coronary artery, with reports of 6-25 dyn/cm^2^.^25-27^ Adherent ECs were re-plated for 3 hrs and subjected to fluid SS again. Control whole cell populations were also plated for 3 hrs. The adherent re-plated ECs exhibited a mean cell retention of 64.1±14.6% compared to 30.4±11.0% (p<0.001) for the whole population (Figure 1F). Because exposing ECs to SS can stimulate focal contact formation and increase adhesion strength,^28^ these observations could be the result of adaptation to SS rather than a distinct subpopulation of adherent cells. To eliminate this possibility, we performed an independent experiment using trypsin as a method to enzymatically detach and sort the cells based on adhesion strength. The non-adherent, adherent and whole population groups were isolated based on detachment kinetics upon incubation with trypsin. Following exposure to fluid SS, the adherent subpopulation exhibited a mean cell retention of 49.2±21.2% compared to 31.4±8.3% for the whole population and 24.3±8.3% for the non-adherent subpopulation (Figure S1). The mean retention of the adherent subpopulation was significantly greater compared to the non-adherent subpopulation (p<0.05). From these experiments, we conclude there exists a distinct subpopulation of ECs with a higher likelihood of resisting detachment under fluid SS *in vitro*. Therefore, cell detachment is not an entirely random process, and it may be possible to predict which ECs will remain adherent when seeded onto a vascular graft.

**Figure 1.**
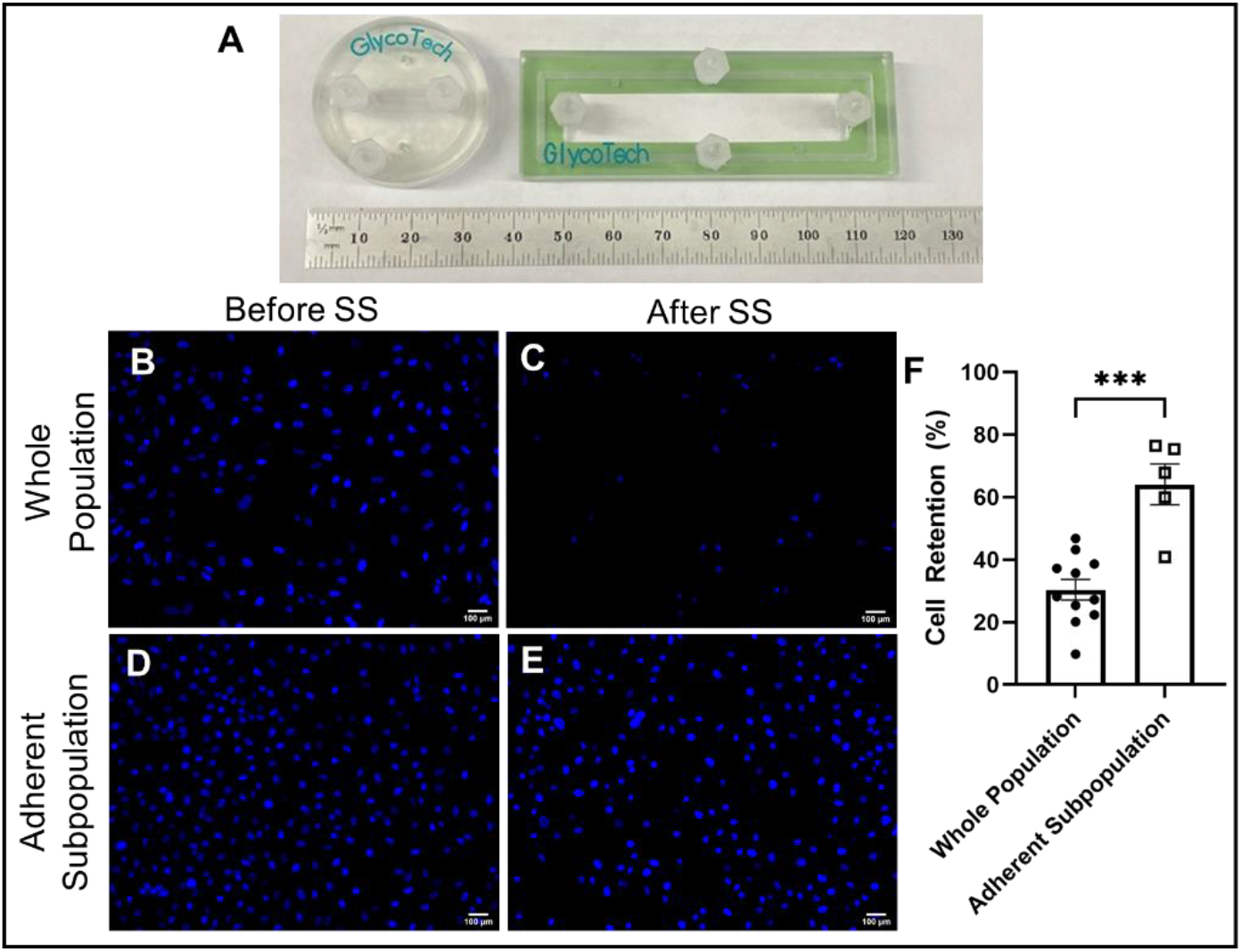
Sorting a population of endothelial cells (ECs) using fluid shear stress (SS) reveals an adherent subpopulation, which has augmented retention when re-exposed to fluid SS (A) Photo of Glycotech parallel plate flow chambers used to subject the ECs to fluid SS (B-E) 10x images of nuclei of the whole population (B,C) and adherent subpopulation (D,E) before and after exposure to SS (F) Adherent subpopulation isolated using fluid SS shows significantly higher cell retention. Unpaired t-test, ***p<0.001. n=5-11 plates per group

### Endothelial cells in the adherent subpopulation show increased vinculin-positive focal adhesions (FAs)

Next, we used vinculin staining to visualize the focal adhesions (FAs) in the cultured whole population and adherent subpopulation ECs (Figure 2). The average number of vinculin-positive FAs normalized to cell number was not significantly different between the groups (122.7±50.4 for the whole population and 127.6±111.6 for the adherent subpopulation) (Figure 2K). However, the total area of the vinculin-positive FAs normalized to cell number was significantly greater in the adherent subpopulation with an average of 0.74±0.40 µm^2^ compared to 0.21±0.09 µm^2^ for the whole population (p<0.05) (Figure 2L). This suggests there are differences in the size of vinculin-containing FAs of the adherent subpopulation, which may contribute to the higher retention observed.

**Figure 2.**
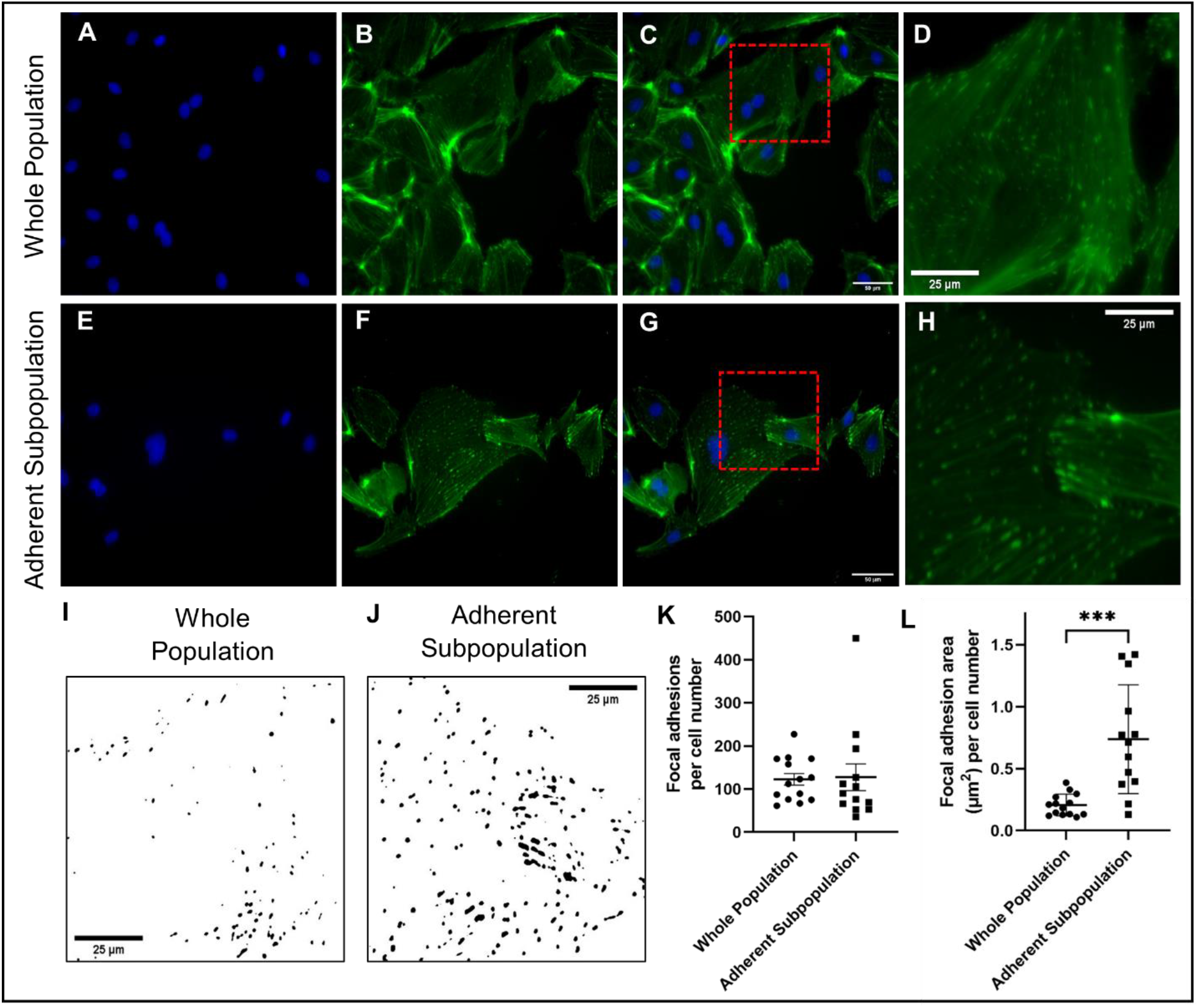
Adherent subpopulation has increased normalized focal adhesion (FA) area. (A-D) Whole population and (E-H) adherent subpopulation representative images for FA analysis using nuclei (blue) and vinculin (green) staining. Analyzed using ImageJ based method described by Horzum et al. 40X images. Red dashed circles in panels C,G shows area zoomed in for panels D,H (I-J) Representative binary images panels D,H to show FAs used for quantification (K) FAs per number of cells in the image did not show a significant difference between the groups (L) FA area per cell number was significantly greater for the adherent subpopulation. Unpaired t-test, ***p<0.001, n=13-14 images from 3 plates per group

### RNA-sequencing identifies differentially expressed genes in the adherent subpopulation

We hypothesized that differential gene expression of molecular signals controlling cell adhesion may contribute to the observation that some ECs are able to resist detachment under exposure to fluid SS. To characterize the transcriptome of the adherent ECs, bulk RNA-seq was used to examine mRNA content of ECs that remained adherent following exposure to 5, 10, or 30 min fluid SS (adherent subpopulations) relative to 0 min fluid SS (whole population) (Figure 3). RNA-seq identified 2017, 1431, and 1255 differentially expressed genes (p<0.05) in adherent ECs isolated after 5, 10, or 30 min of an average 15 dyn/cm^2^ fluid SS, respectively, when compared to the whole population (Figure 3C). Among the differentially expressed genes, there were 156 genes common in the 5, 10, and 30 min adherent subpopulations.

**Figure 3.**
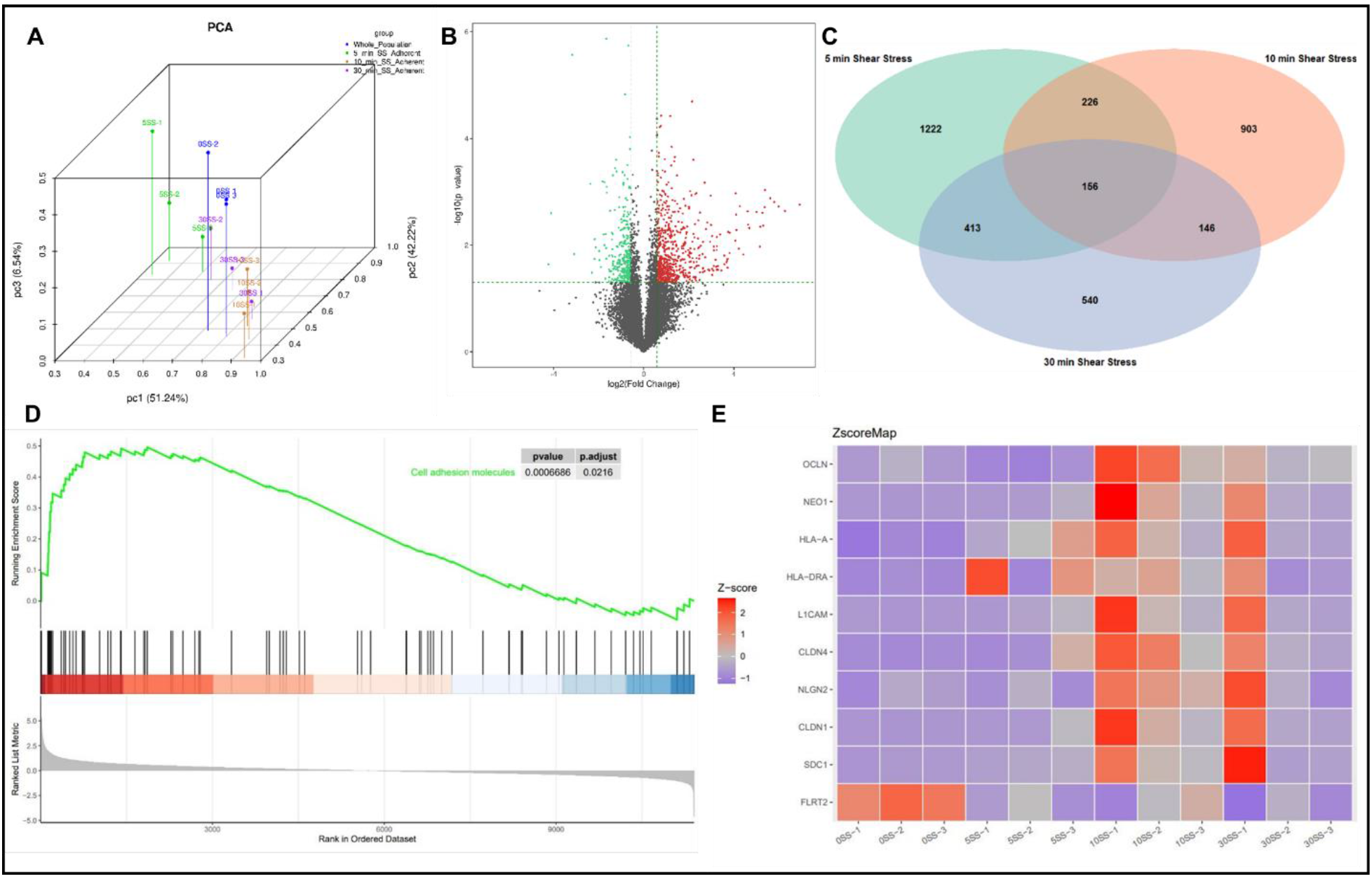
RNA-sequencing identifies the “cell adhesion molecules” pathway is enriched in the adherent subpopulations compared to the whole population (A) Principal component analysis (PCA) plot for all samples (B) Volcano plot for the 10 min adherent subpopulation compared to the whole population. Red dots show upregulated genes and green dots show downregulated genes. Gray dots show genes that were not differentially expressed (C) Venn diagram showing the differentially expressed genes in each adherent subpopulation compared to the whole population (E) Gene set enrichment analysis (GSEA) plot for cell adhesion molecules in the 10 min adherent subpopulation (E) Z-score heatmap of significant genes from the “cell adhesion molecules” pathway

Pathway analysis revealed enrichment in “cell adhesion molecules” for the adherent subpopulation (Figure 3D). Of the 26 genes in the “cell adhesion molecules” pathway, 9 were significantly differentially expressed in the adherent subpopulation isolated after 10 min of SS (Figure 3E). From the 156 differentially expressed genes common to the adherent subpopulations, fibronectin leucine rich transmembrane protein 2 *(FLRT2)* was among the significantly downregulated genes and was explored further based on its expression on the EC surface and existing literature supporting its role in cell adhesion.^29-31^

### Endothelial cells in the adherent subpopulation express low levels of FLRT2

The mRNA data from RNA-seq was first validated with RT-qPCR. Significantly lower *FLRT2* mRNA content was evident in the 5, 10, and 30 min fluid SS adherent samples relative to the 0 min fluid SS samples (Figure 4A). The relative fold change was 0.31±0.07, 0.47±0.09, and 0.45±0.10 for the 5, 10, and 30 min SS samples, respectively, compared to 1.00±0.15 for the 0 min SS. Next, we validated FLRT2 protein content using flow cytometry (Figure 4B). The 0 min fluid SS (whole population) cells showed higher expression of cell surface FLRT2 compared to the 10 min fluid SS (adherent subpopulation) cells, quantified using mean fluorescence intensity of Alexa Fluor 488 (1320.0±63.9 for 0 min SS, 1122.0±26.0 for 10 min SS, 320.2±36.9 for unstained control, and 340.8±14.4 for IgG control, ANOVA p<0.0001) (Figure 4C). Based on the confirmation of cell surface FLRT2 expression differences in the adherent subpopulation compared to the whole population, we explored cell sorting using FACS.

**Figure 4.**
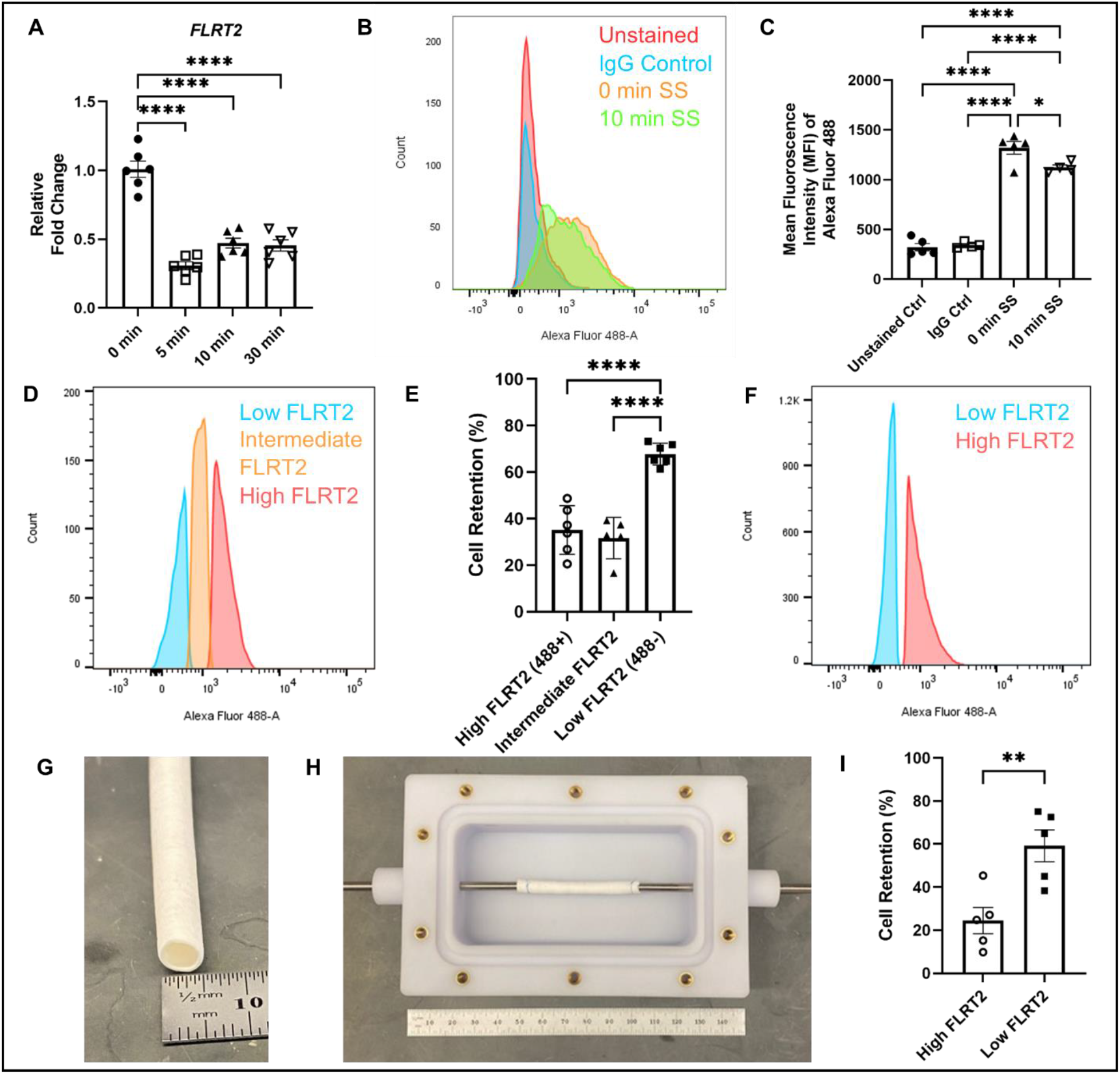
Endothelial cells (ECs) with low FLRT2 expression show increased retention when exposed to fluid shear stress (SS) (A) RT-qPCR results show decreased *FLRT2* mRNA levels in the adherent subpopulations compared to the whole population (B) Histogram of flow cytometry results. Unstained control (red), IgG control (blue), 0 min fluid SS whole population (orange), 10 min fluid SS adherent subpopulation (green) (C) Quantification of mean fluorescence intensity (MFI) of Alexa Fluor 488 for FLRT2 staining with flow cytometry (D) Histogram of gating into low, intermediate, and high FLRT2 expression groups with fluorescence activated cell sorting (FACS) (E) Cell retention (%) for ECs expressing low, intermediate, or high levels of FLRT2 on TC plates after 30 min of 15 dyn/cm^2^ fluid SS (F) Histogram of gating into low and high FLRT2 expression groups with FACS for retention on vascular graft experiments (G) Image of the 4 mm inner diameter graft (H) Image of a vascular graft sample mounted in the perfusion chamber (I) Cell retention (%) for ECs expressing low or high levels on vascular grafts after 30 min of 10 dyn/cm^2^ fluid SS. ANOVA with Tukey’s analysis. * p<0.05, ** p<0.001, **** p<0.0001. N=5-6 samples per group

### Endothelial cells sorted for low FLRT2 expression show enhanced retention when exposed to fluid shear stress

FACS was used to sort primary pooled HUVECs into low, intermediate, and high FLRT2 expression groups for EC retention studies (Figure 4D). The goal was to identify cell populations with high, intermediate, and low likelihood of retention upon exposure to fluid SS, respectively. The sorted HUVECs were cultured overnight and subjected to 15 dyn/cm^2^ fluid SS with a parallel plate flow chamber for 30 min. The retention of the ECs sorted for low surface expression of FLRT2 was 67.7±4.3%, which was significantly greater than 35.1±9.5% for the high and 31.6±8.0% for the intermediate groups (ANOVA, p<0.0001) (Figure 4E). Based on the similar retention between the high and intermediate FLRT2 surface expression groups, ECs were sorted into low and high groups for experiments on vascular grafts (Figure 4F). Sorted ECs were cultured on fibronectin-coated grafts overnight. The grafts were exposed to an average fluid SS of 10 dyn/cm^2^ with a pulsatile pump for 30 min in a custom chamber (Figure 4H). This SS value was chosen based on the average wall SS in the coronary artery, which is around 7 dyn/cm^2^,^25^ and the flow capabilities of the pulsatile blood pump to achieve the required 405 mL/min flow rate. The retention of the ECs sorted for low FLRT2 was 59.2±7.4%, which was significantly greater than the retention of 24.5±6.1% for ECs sorted for high FLRT2 (t-test, p<0.001) (Figure 4I). Therefore, ECs with low FLRT2 expression show enhanced retention on vascular grafts exposed to fluid SS.

## Discussion

Here we report that EC detachment upon exposure to fluid SS is not an entirely random process, and we can select for the ECs in a population that are more likely to remain adherent. We show that ECs can be sorted based on FLRT2 expression, and the ECs expressing lower levels of FLRT2 have enhanced retention under exposure to fluid SS *in vitro*.

There is a strong clinical need to develop a technique to routinely endothelialize blood-contacting devices.^32-35^ The critical barrier is poor retention of ECs on graft biomaterials upon exposure to physiologic flow conditions.^10,36,37^ Our method of sorting ECs based on FLRT2 expression is an innovative and straightforward approach that selects for the ECs more likely to remain adherent at the time of seeding. Cell-sorting with FACS provides a safer, cheaper option than cell modification technologies such as CRISPR-based gene editing.^38^ The sorted highly adherent ECs resist detachment when exposed to fluid SS and show promise for future applications on blood-contacting medical devices.

When choosing a molecular signal from the RNA-seq data, we sought to identify a differentially expressed gene that was common to all the adherent subpopulations and would be conserved across different EC types and species. We were not interested in mechanosensitive genes specifically, but also did not rule out those genes as potential options. The list of common differentially expressed genes included ones that were significantly upregulated and downregulated in the adherent subpopulations compared to the whole population. Genes that encode EC surface proteins were of particular interest since those could be targeted with an antibody while maintaining cell viability. This would allow a population of ECs to be sorted to select for the highly adherent subpopulation. We explored FLRT2 as a possible target based on the significant downregulation of *FLRT2* gene expression in all adherent subpopulations and existing literature.

Fibronectin leucine-rich transmembrane proteins are a family of widely expressed cell-surface molecules,^39^ and FLRT2 is expressed on HUVECs. FLRT2 has been previously reported to have both repulsive and adhesive functions.^29,30^ However, the role of FLRT2 in EC retention after exposure to SS has not been investigated previously.

The mechanism by which FLRT2 downregulation allows for enhanced retention of ECs exposed to SS is still largely unknown. Our study showed that we can selectively sort for the more adherent ECs in a population based on low FLRT2 expression, which was the goal of the study. The quantification of vinculin-containing FA number and area indicates that the size of the vinculin-containing FAs in the highly adherent ECs is greater, which could suggest the mechanism controlling EC detachment is related to FA formation. This aligns with observations from Camilo et al., which reported that silencing FLRT2 expression in ECs increases the number and size of vinculin-containing FAs.^31^ Camilo et al. concluded that FLRT2 is an endogenous ligand that activates latrophilin-2 (LPHN2) and promotes the disassembly of integrin-based FAs. Other published literature on FLRT2 by Ando et al. described how deletion of *FLRT2* in mice leads to an increased number of mature vessels in a tumor cancer model.^30^ Silencing *FLRT2* with siRNA caused downregulation of cytoskeletal genes (*RHOB, RASA1, ROCK1*) and cell cycle genes (*CDC45, E2F8*) and upregulation of intercellular adhesion genes *(CEACAM1, LGALS9*). Interestingly, we also saw downregulation of *RHOB & E2F8* and upregulation of *LGALS9* in the adherent subpopulations compared to the whole population RNA-seq results, which further supports the notion that *FLRT2* expression is having an effect in the adherent subpopulation. However, the mechanism through which low FLRT2 expression improves the retention of ECs exposed to fluid SS warrants further investigation.

We confirmed that FLRT2 protein content matched the mRNA data using flow cytometry. Interestingly, the flow cytometry analysis of the HUVECs showed a wide range of FLRT2 expression intensity between individual cells. This may be due to the heterogeneity of FLRT2 expression in cultured HUVECs, which could explain the adherent and non-adherent phenotype ECs. The reason for the heterogeneity is unknown and warrants further investigation.

For clinical translatability, it is ideal to have an autologous EC source for seeding the small-diameter vascular grafts.^40^ BOECs, which exhibit the hallmarks of mature ECs, can be generated from the circulating peripheral blood mononuclear cell populations. This makes them a promising autologous EC source for adult patients, without a need for immune suppressing drugs.^13^ Previous work has described the suitability of BOECs for TEVG applications^5,41^ and similarity of gene expression responses to SS in human BOECs (HBOECs) compared to HUVECs.^42^ Further, the global gene expression profile of HUVECs, endothelial colony-forming cells, and human coronary artery endothelial cells is known to be similar.^43^ This provides evidence that our findings with highly adherent HUVECs can be translated to other EC populations, including BOECs. For future preclinical studies in a porcine model, we plan to seed grafts with autologous porcine BOECs that have been enriched for the adherent subpopulation.

One limitation of our study relates to our approach of using 0, 5, 10, and 30 min fluid SS groups aimed to identify a molecular signal that was consistent among the adherent subpopulations and possibly trending with duration of retention. It is well known that ECs are highly sensitive to SS and changes in gene expression can be observed within 40 min after exposure to SS.^44^ Short exposure times to fluid SS were chosen to isolate the adherent HUVECs in order to minimize confounding effects of the fluid SS response. The goal was to identify molecular signals unique to the adherent ECs, not to study the molecular signaling response to SS. *FLRT2* has not been previously reported as a mechanosensitive gene,^45^ but it is difficult to select for cells able to resist detachment upon exposure to fluid SS without simultaneously activating cell response to SS.

Another limitation is the retention of low FLRT2-expressing HUVECs was only around 63%, meaning 37% of cells were still lost. Although this is a substantial improvement from ∼30% retention, it is currently unknown what percentage of cell retention is necessary to achieve long-term graft patency. The biological response including thrombosis and neointimal hyperplasia needs to be studied in an animal model. If higher retention is necessary, the results of this study allow for numerous approaches to be investigated. For example, a larger starting population of cells can be used with a more stringent sorting strategy to select for cells with very low expression of FLRT2. In addition, the RNA-seq data provided several candidate molecular signals that can be used to design sorting strategies based on the expression level of multiple genes.

We have shown for the first time that a population of ECs can be sorted to select for a subpopulation of highly adherent ECs which resists detachment when exposed to fluid SS. This is a promising approach for improving EC retention on a vascular graft and future work will investigate retention of sorted ECs *in vivo* using a pre-clinical large animal model. This represents an innovative method to seed a small-diameter vascular graft with ECs that are more likely to remain adherent upon exposure to circulating blood.

## Acknowledgments

We kindly acknowledge Christopher Monti for assistance with the RT-qPCR experiments, the Engineering Core at the Medical College of Wisconsin for fabricating the graft perfusion chambers, Toby Frost for helping test the perfusion chambers, the flow cytometry core at the Versiti Blood Research Institute for assistance with the FACS experiments, and Ab Medica S.p.A. for donating the DEGRAPOL® DP15.

